# Effect of earthworms in removal and fate of antibiotic resistant bacteria (ARB) and antibiotic resistant genes (ARG) during clinical laboratory wastewater treatment by vermifiltration

**DOI:** 10.1101/2020.10.01.321885

**Authors:** Sudipti Arora, Sakshi Saraswat, Ankur Rajpal, Harshita Shringi, Rinki Mishra, Jasmine Sethi, Jayana Rajvanshi, Aditi Nag, Sonika Saxena, A.A. Kazmi

## Abstract

The wastewater treatment plants effluent has been implicated in the spread of antibiotic resistant bacteria (ARB) as these environment contains multiple selective pressures that may increase mutation rates, pathogen survivability, and induce gene transfer between bacteria. In lieu of this, the present study explored the dynamics of earthworm-microorganisms interactions on the treatment efficacy of clinical laboratory wastewater treatment by vermifiltration and the effect of earthworms in the fate of removal of pathogens and ARB. The results of the study showed that earthworms and VF associated microbial community had a significant effect on BOD and COD reduction (78-85%), pathogen removal (>99.9 %) and caused a significant shift in the prevalence pattern of ARB. Additionally, molecular profiling of ESBL (bla_SHV_, bla_TEM_ and bla_CTX-M_), MRSA (*mec-A)* and Colistin (*mcr*-1) gene confirmed the probable mechanisms behind the resistance pattern. The microbial community diversity assists in the formation of biofilm, which helps in the removal of pathogens and results in a paradigm shift in the resistance profile of ARB and ARG, specifically most effective against drugs, targeting cell wall and protein synthesis inhibition like Ampicillin, Ticarcillin, Gentamicin and Chloramphenicol. These findings prove vermifiltration technology as a sustainable and natural treatment technology for clinical laboratory wastewater.

## 1. Introduction

In the recent years there has been growing concern about the release of organic compounds of anthropogenic origin, known as emerging organic contaminants and xenobiotics such as antibiotics, to the environment. The presence of antibiotics is of special concern due to the development of antibiotic resistant bacteria (ARB) and antibiotic resistance genes (ARGs). These substances are extensively used in both human and veterinary medicine against microbial infections and are excreted from the body of the treated organisms, together with their metabolites, within a few days of consumption. Recent studies have shown that incomplete metabolism in humans and improper disposal of antibiotics to wastewater treatment plants (WWTPs) has been a main source of antibiotic release into the environment [1]. It has been widely demonstrated that conventional WWTPs are inefficient for the removal of many antibiotics, ARBs and ARGs, thus contaminating the receiving ecosystems with a complex mixture of bioactive agents and bacteria. It is estimated that 30–60% of all prescribed antibiotics can end up in WWTPs, which act as primary reactors creating ARB and ARGs [2]. Once in the environment, antibiotics can lead to the continuous selection for ARB that contains ARGs. In urban areas, the regular discharges of WWTP effluents into aquatic bodies (e.g. rivers, lakes), including hospital wastewater effluents, is the main entrance pathway of these substances into the environment. Over the years, WWTPs have been recognized as hotspots for ARB which enters the environment. When these raw and conjugated metabolites reach the WWTPs and finally to the groundwater, rivers, lakes, oceans and soil, it can be very harmful to aquatic organisms and form ARBs and ARGs [3]. Due to the global rise in antibiotic resistance, additional emphasis is now being placed on the detection and removal of pharmaceuticals, ARB and ARGs, and pathogens from treated wastewater but federal and state monitoring programs for these pollutants remains minimal or nonexistent [4]. It has been reported that estrogens can cause hermaphroditic fish, while some painkillers are poisonous to trout and certain psychopharmaceuticals can affect fish and bird behavior [5]. Thus, it is highly recommended to perform preliminary onsite treatment for these effluents. Despite the constant improvements being made over past decades in the field of sanitation & wastewater treatment, unsafe management of fecal waste and wastewater still presents a major risk to public health and the environment. This problem is multifold by the discharge and addition of hospital and clinical laboratory wastewater into municipal WWTPs. Hospital and clinical wastewater is quite different from the wastewater discharged from other sources and is hazardous and infectious. It consists of a wide range of several micro and macro-pollutants, discharged from operation (surgery) rooms, wards, laboratories, laundry, polyclinics, research units, radiology, and medicine and nutrient solutions used in microbiology and diagnostics laboratories [6]. Regardless of the stringent guidelines issued by concerned authorities for onsite treatment of hospital & clinical wastewater (WHO [7], CPCB [8]) the hospitals of the majority of developing countries discharge their effluent directly to the public sewer network without any prior treatment. Since, municipal WWTPs are not designed to deal with these kind of biological waste, as a result, pathogens, microorganisms and antibiotics are known to survive in the urban wastewater treatment plants and, subsequently, in the treated effluent [9]. As a compensation process against antibiotics, bacteria present in wastewater generate antibiotic resistance by mutations or by horizontal gene transfer [10,11]. Horizontal gene transfer is made possible in large part by the existence of mobile genetic elements (i.e., transposons and integron-associated gene cassettes) that can efficiently contribute to the acquisition, maintenance and spread of ARGs within bacterial communities [12].

In spite of this hazardous influence, there are only few studies investigating the release and direct impact of hospitals and clinical laboratory wastewater into the environment or communal sewage system. In WWTPs receiving hospital sewage, this effect may be more pronounced due to increased concentrations of antibiotics, ARB, and clinical pathogens entering the treatment system from hospital sewage [4]. As the number of resistant bacterial infections continues to rise worldwide, wastewater treatment processes that focus on the removal of antibiotics of critical importance for human health, and is necessary to reduce the selection and spread of bacteria resistant to these antibiotics.

In addition to this, clinical wastewater contains various types of ARGs, including those resistant to antibiotics such as tetracyclines, β-lactams, sulphonamides, macrolides, fluoroquinolones, and even multiple drug resistance [13,14]. Different types of ARGs confer microbial resistance to different antibiotics [15,16]. Centralized technologies like membrane bioreactor combined with activated sludge process are gaining wide acceptance for hospital and clinical laboratory wastewater treatment due to its high removal capacity for bacteria and antibiotics. However, installation, operation, and maintenance cost form a major challenge towards the establishment of these technologies [3]. Thus, it is imperative to develop and utilize an efficient and cost-effective technology.

In the recent years, earthworms driven wastewater treatment process, known as vermifiltration technology has shown tremendous benefits and proven successful in treating domestic wastewater [17,18], industrial wastewater like herbal pharmaceutical wastewater [19], gelatine wastewater [20], dairy wastewater [21], etc. The present study investigates the impact and efficiency of earthworms in contaminants removal, pathogens and ARB removal of an on-site vermifilter for treating clinical laboratory wastewater. The effect of earthworms in treating clinical laboratory wastewater and their impact on ARB and ARG occurrence and further concentrations in influent and effluent are compared to identify potential differences. A clinical diagnostic laboratory deals with urine, blood and faecal samples of infected patients, there are the chances of detection of antibiotics present in their samples, and it is imperative to know the resistance pattern of the microorganisms present in the influent and effluent. In view of this, the present study was carried out to understand the mechanistic insights into vermifiltration technology for clinical wastewater treatment. The main objective of the present study is to determine the performance efficacy of vermifilter and to understand the mechanistic insights by deciphering the role of earthworms in the treatment process. This study also aimed to construe the role of earthworms in AMR and ARB reduction. To the best of our knowledge, this study reports for the first time, the efficacy of the vermifiltration process for the treatment of clinical laboratory wastewater and its effect on antibiotic resistance.

## 2. Materials and Methods

### 2.1. Experimental design

The study was performed at Dr. B. Lal Institute of Biotechnology, Jaipur, India. A pilot-scale vermifilter (VF) of 1 KLD treatment capacity (material - concrete) with dimensions of 1 m^3^ was installed at Dr. B. Lal Clinical Laboratory Pvt. Ltd. Campus, Jaipur, in field conditions for treating clinical laboratory wastewater over a period of one year (Jan 2019-Dec 2019). The wastewater was collected by a pump to a collection tank, which was then transferred to a pretreatment or overhead tank, from where the wastewater flowed to the VF by gravity and is then connected to an effluent collection tank The pretreatment tank contained a mixer for constant mixing of wastewater and sludge and a static ball for controlling water level. The wastewater distribution system in the VF consisted of polyvinyl chloride (PVC) perforated pipes. The working principle of the VF, along with all the design parameters is shown in Figure 1, adopting the methodology as described by Arora et al., 2014 [17]. The VF comprised of a filter bed consisting of different media. It was made up of 3 layers of river bed materials, including a layer of coarse size gravel (aggregate size 12-14 mm, 15 cm thickness) as bottom supporting layer, a layer of medium size gravel (aggregate size 6–8 mm, 15 cm thickness) and a layer of fine gravel (aggregate size 2-4 mm, 15 cm thickness) at the top. At the top, VF was filled with 30 cm fine vermigratings and cow dung (0.05–5 mm), as the active layer and empty space (15 cm) is kept at the top for aeration purpose. The active layer containing vermigratings had a uniformity coefficient of 1.36, the effective size of 0.118 mm and a density of 1517.6 kg/m^3^. It had an average pH of 6.5-7.0, a density of 96 kg/m^3^ containing lignocellulosic material and the VF rarely produced odor during its long-term operation period, which showed that the VF was aerobic throughout the process. The VF was inoculated with earthworm’s species *Eisenia fetida* with stocking density of 10,000 worms/m^3^. The clinical laboratory wastewater was allowed to flow to the VF with a hydraulic loading rate (HLR) of 1.0 m^3^/m^2^/day for 12 hours continuously and hydraulic retention time (HRT) came out to be of 7-8 hours. The temperature in the VF bed was monitored under standard conditions, with temperature maintained throughout the study period, ranging between 20-30°C.

**Figure 1:**
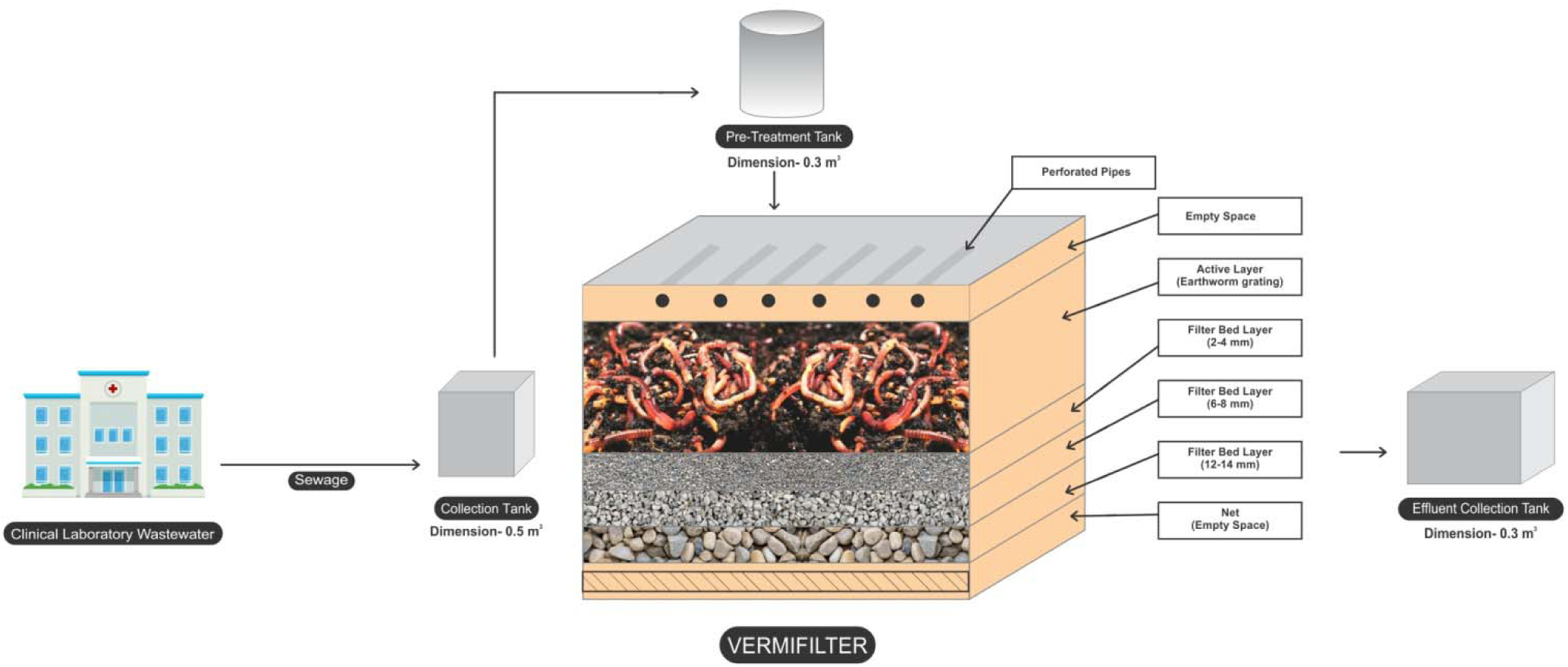
Schematic flow diagram of vermifilter for treatment of clinical laboratory wastewater

### 2.2. Performance evaluation of vermifilter

The performance evaluation of vermifilter was determined by physicochemical and microbiological parameter analysis of influent and effluent samples. Sampling was performed once a week for a period of one year. The physical parameter included pH, temperature, and electrical conductivity (EC). The chemical parameter included dissolved oxygen (DO), total dissolved solids (TDS), salinity, biochemical oxygen demand (BOD) and chemical oxygen demand (COD), and nutrients (ammonium (NH_4_^+^) and nitrate (NO_3_^−^)). The microbiological parameter included total coliforms (TC), fecal coliforms (FC), fecal streptococci (FS), *E. coli* and *Salmonella*. All the parameters were analyzed as per the standard methodology by Arora et al., 2020 [22]. The analysis was performed on the same day of sampling and when the same-day analysis was not possible, samples were stored at 4°C for less than 24 hours in sterile container bottles which were screwed tightly to avoid external contamination.

### 2.3 Investigation of microbial community diversity

The microbial community diversity inside the VF was investigated by culture based method. The spread plate technique was used to isolate microorganisms from the influent, effluent, earthworm gut, coelomic fluid (CF) and filter media of the VF. Sampling was done twice during the operation period. The bacterial colonies were isolated on nutrient agar media and incubated at 30°C for 24 h. Well-isolated colonies were selected and sub cultured and cells from the new colonies were then picked up with an inoculating needle and transferred onto agar slants to be maintained as pure culture. Pure bacterial cultures were then identified and classified morphologically (shape, gram staining and motility) and biochemically (sugar utilization, indole production, citrate utilization, methyl red-Voges Proskauer (MR-VP), triple sugar iron (TSI) utilization, oxidase production, catalase production, gas production from glucose and coagulase test) according to Bergey’s Manual [23].

### 2.4. Determination of antibiotic susceptibility using double disk synergy test

Antibiotic susceptibility test was done by disk diffusion method [24]. The isolates obtained from influent and effluent were swabbed onto the surface of Mueller Hinton Agar (MHA) media plate (diameter-150 mm). The filter paper disks impregnated with a standardized concentration of antibiotics were placed on the surface, and the size of the zone of inhibition around the disk was measured after overnight incubation at 35°C. The 12 antibiotic disks used are mentioned in Table 1. These are further categorized upon the basis of their class and the mechanism involved in their antibacterial action. Each of these antibiotics have their own specific mechanism and they were maintained according to the guidelines outlined in the Clinical and Laboratory Standards Institute [CLSI] documents [25].

**Table 1:** List of antibiotics used for the detection of ARB along with their class and mechanism of action

| Antibiotics | Class of Antibiotics | Mechanism of Action |
| --- | --- | --- |
| Amoxicillin (AMX<sup>10</sup>) | Penicillin | Inhibition of cell wall |
| Ampicillin (AMP<sup>10</sup>) |  |  |
| Ticarcillin (TCC<sup>75/10</sup>) |  |  |
| Ceftazidime (CAZ<sup>30</sup>) | Cephalosporin |  |
| Cefotaxime (CTX<sup>10</sup>) |  |  |
| Ceftriaxone (CTR<sup>10</sup>) |  |  |
| Streptomycin (S<sup>10</sup>) | Aminoglycoside | Protein synthesis inhibitors |
| Gentamicin (HLG<sup>120</sup> ) |  |  |
| Erythromycin (E<sup>5</sup>) | Macrolide |  |
| Tetracyclines (TE<sup>10</sup>) | Tetracycline |  |
| Chloramphenicol (C<sup>10</sup>) | Chloramphenicol |  |
| Ciprofloxacin (CIP<sup>10</sup>) | Fluoroquinolone | Inhibition of nucleic acid |

### 2.5. Molecular profiling of ARB genes

To further understand the underlying molecular mechanism, identification of the production of ESBL (extended spectrum beta-lactamases), MRSA (methicillin-resistant *Staphylococcus aureus*) and Colistin genes was performed using culture independent method [26]. Metagenomic DNA was extracted using the Power water DNA Isolation Kit (Qiagen) following the manufacturer’s protocol with modifications. This kit is specifically designed for isolating DNA from filtered environmental water samples and includes inhibitor removal technology aimed at removing humic acid and other organic matter commonly found in environmental samples that can interfere with downstream analyses. The protocol modifications included an incubation at 65°C for 10 minutes following addition of solution P1 and a lysozyme digestion (final concentration = 10 mg/ml) at 37°C for 1 hour and an RNase digestion at 37°C for 20 minutes immediately following the bead-beating step. DNA quantity and quality were estimated using Nanodrop-1000 Spectrophotometer (Nanodrop Eppendrof). DNA concentrations ranged from 75 to 130 ng/μl per sample. The presence of bla_TEM_, bla_SHV_, bla_CTX-M_ (ESBL), mec-A (MRSA) and mcr-1 (Colistin) genes in transformants were confirmed by PCR. Approximately 2-3 μg of total DNA for each sample was used for a PCR run. Primers for genotyping were obtained from Eurofins by custom order. The primer pair used for amplification consisted of 27F (5′-AGA GTT TGA TC[A/C] TGG CTC AG-3′) and 1492R (5′-G[C/T]T ACC TTG TTA CGA CTT-3′).

## 3. Results and Discussions

### 3.1 Performance efficacy of VF by physicochemical and microbiological parameters analysis

The performance efficiency was checked thoroughly by analysis of physicochemical and microbiological parameters. For all the physicochemical parameters carried out over a period of 12 months, the average value was calculated and it has been depicted in Figures 2 and 3. Radical variations were observed in the pH of the influent ranging from 4.2 to 9.3 as clinical wastewater contains a variety of toxic or persistent substances such as pharmaceuticals, radionuclides, chemical hazardous substances, pathogens, radioisotopes, solvents, endocrine disruptors, detergents, radiographic developers, and disinfectants for medical purposes in a wide range of concentrations. Despite this, it was gradually neutralized to the range of 6.7-7.0 in the effluent referring to the avid buffering capacity of the earthworms (Figure 2a) [21]. Temperature is another key factor that affects earthworm but it was found to be in the range of 20-30°C (Figure 2b) even during extreme summers and winters and didn’t show any harsh effect on the earthworms. This consistency was mainly because of the material used for the construction of VF tank and river bed materials as filter media. The average EC in the influent and effluent seemed lie in the range of 2.9-3.1 mS/cm (Figure 2c). No significant difference was observed between the average salinity of the influent and effluent throughout the study as shown in Figure 2d. During the vermifiltration process, TDS concentration in VF effluent was reduced by 30% from the influent (Figure 3a). This may be attributed to the ingestion of organic and inorganic solid particles in wastewater by earthworms, which excrete them as finer particles. These finer particles are further trapped in the voids of VF and cause high removal efficiency of TDS from wastewater. This is the reason why VF does not choke throughout the operation period but rather work smoothly and uninterruptedly which makes it an excellent choice [27]. TDS purification and removal rates were consistent during all the months as the main processes involved are filtration and sedimentation. DO is another important parameter and the mean DO concentration in the influent was 0.53 mg/L. In VF effluent, the mean DO was observed to be 3.1 mg/L (Figure 3b). DO signify the environmental conditions prevailing inside the VF and it is one of the significant parameters in outflow because low oxygen level is toxic to earthworms and prevailing aerobic microorganisms. The increase in DO is also attributed to the dropping overflow of the VF layers, as water flowed through it naturally by gravity and earthworm also provide aeration through their burrowing activity in the filter bed [28]. Higher DO in the effluent reduces the septic condition and brings a good chance of wastewater reuse for irrigation purposes [29]. DO content is also closely related to organic matter removal calculated in terms of BOD and COD. These are two most essential parameters which define the profile of any type of water. The average BOD in the influent that was observed to be in the range of 200-250 mg/L, which was reduced to less than 30 mg/L in the effluent after the acclimatization period (Figure 3c) whereas the COD was found to be around 390-420 mg/L in the influent decreased to 60-90 mg/L in the effluent (Figure 3d). These results show decrement by 80-85%% and 75-80% for BOD and COD respectively. The reason for high BOD and COD removal is the reduction in organic load present in the influent, by the enzymes produced by the earthworms and microorganisms such as cellulose, amylase and protease. Earthworms also have an ability to enhance the biodegradation of organic matter through their feeding, burrowing and casting behavior. They host millions of decomposer microbes in their gut and excrete them along with nutrients like nitrogen in their excreta. Nitrogen is one of the predominant contaminants in sewage, and nitrates and ammonia result in water pollution and Eutrophication [30]. The average concentration of NO^−^_3_−N in influent was 5.67 mg/L which experienced an increment of 62% in the effluent (Figure 3e). Similarly, NH^+^_3_−N in the influent were detected to be in the range of 6.6-7.0 mg/L while that of the effluent increased to 18-19 mg/L (Figure 3f). This could be attributed to the rapid growth of ammonia oxidizing bacteria at a temperature above 15°C [31]. NH^+^_3_−N was subsequently converted to NO^−^_3_−N via biological nitrification, which was carried out by aerobic autotrophic bacteria using molecular oxygen as an electron acceptor [28]. Meanwhile, high surface DO concentration was beneficial for aerobic microbial survival, which in turn was advantageous for nitrification. All these parameters collectively result in a higher treatment efficiency of the VF.

**Figure 2:**
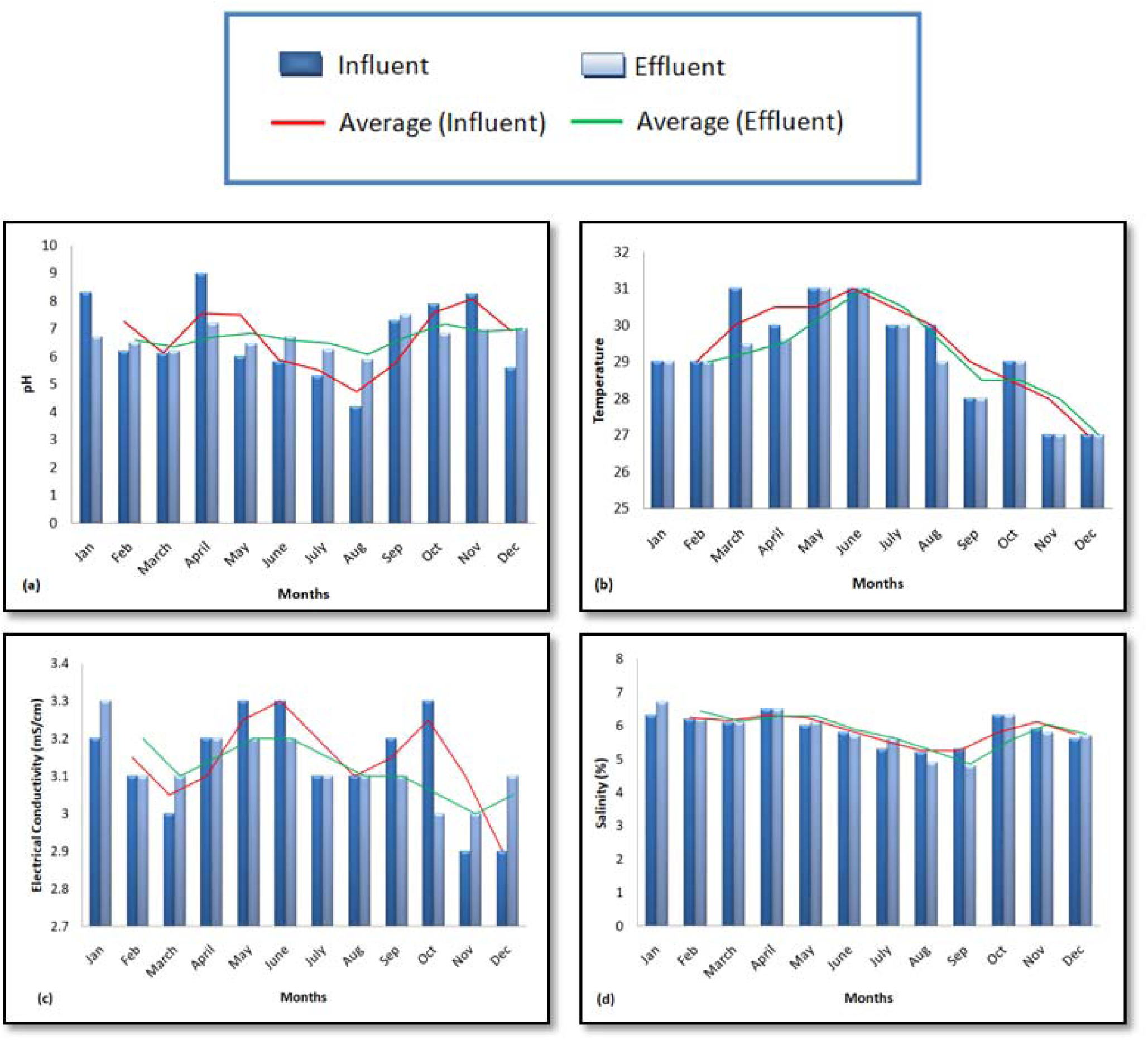
Variations in (a) pH, (b) temperature, (c) EC and (d) salinity over a period of one year.

**Figure 3:**
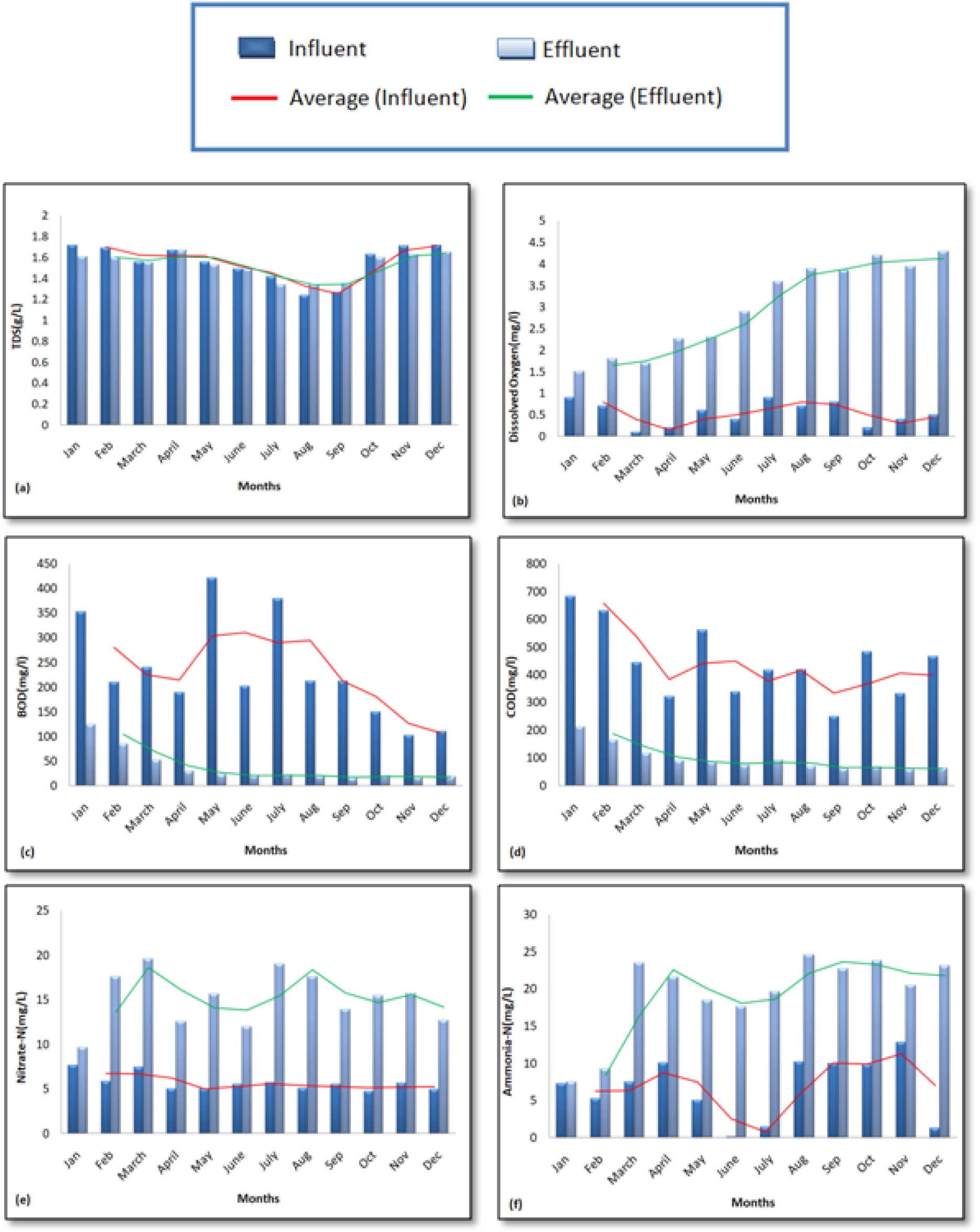
Variations in (a) TDS (b) DO (c) BOD (d)COD (e) Nitrate-N (f) Ammonia-N over a period of one year.

The presence of coliforms is often used as an indicator of overall pathogenicity of the sample. The coliforms and pathogen removal efficacy for wastewater is given in Tables 2. It was observed that the concentration of pathogens in VF was reduced considerably. Log removal values were determined by taking the logarithm of the ratio of pathogen concentration in the influent and effluent and by utilizing these values percentage removal was obtained. The average values of TC, FC and FS were reduced by 90%, 99% and 99% in the effluent, respectively, showing pathogen reduction and quality enhancement of the effluent. The log removal of *E. coli* and *Salmonella* in VF effluent was observed to be 3 with the percentage removal of 99.9%. High removal values may be attributed to the release of coelomic fluid from the body cavity of earthworms as reported by Arora et al., 2014 [17], shows antibacterial properties that inhibit the growth of pathogens. Vermifiltration has shown to reduce pathogens to safe levels through the action of earthworm–microorganisms interactions and antibacterial activity of microflora associated with earthworms [27].

**Table 2:** Changes in microbiological parameters of clinical wastewater during vermifiltration

| Parameters | Influent | Effluent | Percentage Removal |
| --- | --- | --- | --- |
| TC (MPN/100 mL) | 6.7±0.7 | 5.5±0.9 | 90% |
| FC (MPN/100 mL) | 4.5±0.5 | 2.5±0.5 | 99% |
| FS (MPN/100 mL) | 4.3±0.3 | 2.80±0.1 | 99% |
| *E.coli* (CFU/ml) | 3.5±0.5 | 1.2±0.3 | 99% |
| *Salmonella* (CFU/ml) | 3.8±0.5 | 1.67±0.92 | 99% |

### 3.2 Microbial community diversity associated with earthworm-microorganisms interactions

The results revealed the microbial community diversity, based on culture assay showed presence of sixty isolates from influent, effluent, earthworm coelomic fluid, gut of earthworms and filter media layer samples as shown in Figure 4. Out of 60 isolates, 22% belong to the *Micrococcus* sp. (13 isolates), 17% belong to *Bacillus* sp.(10 isolates), 11% belong to *Corynebacterium* sp. (7 isolates), 10% belong to *Proteus* sp., *Pseudomonas* sp. and *Staphylococcus* sp. (6 isolates each), 5% belong to *Shigella* and *E*.*coli* each (3 isolates each), 3% belong to *Citrobacter* sp. and *Klebsiella* sp. (2 isolates each), and 2% belong to *Enterobacter* and *Providencia* (1 isolate each). All these microorganisms occur in the wide range of aerobic to facultative anaerobic bacteria. These results are analogous to the previous studies and contribute for efficient working of a VF system [27,28]. Clinical wastewater outlines the source of various pathogenic and opportunistic pathogens in the influent like *Proteus, Shigella, Pseudomonas, Bacillus* sp. and *Klebsiella* spp. are amongst the most common causes of a variety of community-acquired and hospital-acquired infections [32]. *E. coli* strains are part of the normal microbiota of the gut and are expelled into the environment within fecal matter. *Corynebacterium* sp. acts as pathogens in immune suppressed patients. Interestingly, filter media layer and coelomic fluid was primarily inhabited by gram-positive microflora. On the contrary, isolates obtained from the earthworm gut belonged to the gram-negative microflora. Earthworms are reported to stimulate and accelerate microbial activity by increasing the population of microorganisms and through improved aeration by burrowing actions [33]. The selected isolates obtained from influent and effluent samples were subjected further to antibiotic susceptibility testing.

**Figure 4:**
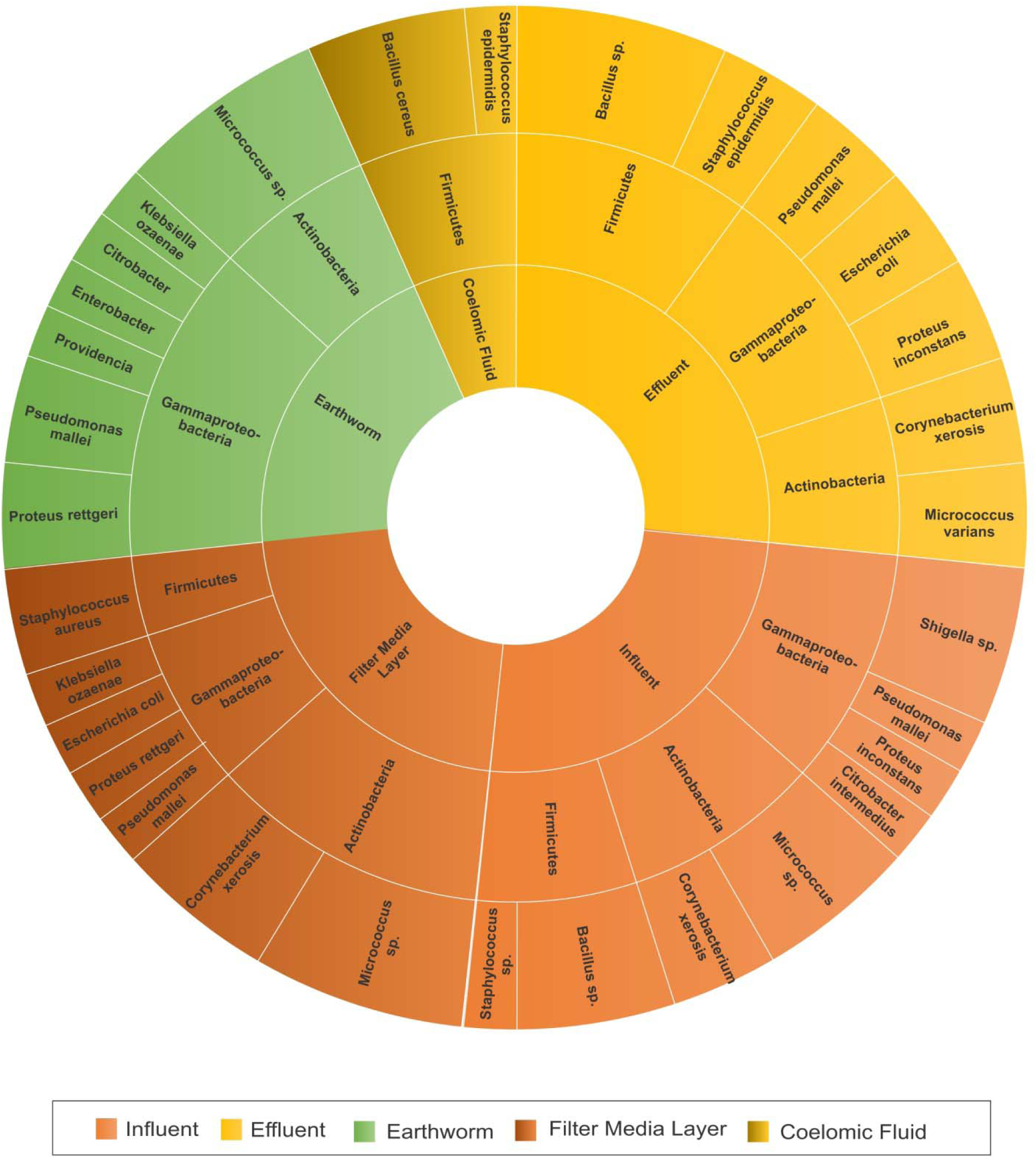
Microbial community diversity isolated from influent, effluent, earthworm gut, coelomic fluid and filter media layer

### 3.3 Antibiotic susceptibility testing

The dissemination of ARB in the environment is a major concern, and WWTPs have been implicated as both reservoirs and contributors of ARB in surface waters following wastewater treatment. We identified the culturable aerobic microbial community composition of influent and effluent that were resistant to antibiotics of critical importance. A total of 12 strains isolated from influent and effluent were selected to determine antibiotic susceptibility test against the drugs as shown in Figure 5. The results indicated that none of the species showed resistance against Ciprofloxacin (CIP10/5) in both influent and effluent. Few of the isolates showed equal resistance in the influent and effluent samples against Amoxicillin (AMX10), Cefotaxime (CTX10), Erythromycin (E5) and Tetracyclines (TE10). Interestingly, few species which were found to be resistant in the influent against Ampicillin (AMP10), Ticarcillin (TCC75/10), Gentamicin (HLG120/ GEN10) and Chloramphenicol (C10) experienced a decrement in the percentage of resistant isolates in the effluent samples. Another intriguing observation was the presence of an isolate which was resistant against Streptomycin (S10/ HLS300) in the effluent while no resistant isolates were observed in the influent. The probable reason for this can either be residual contaminants during the collection or presence of non-culturable bacteria. Thus, to determine the source of resistance and to elucidate the mechanism associated with it, further profiling of ARG was conducted for the effluent sample.

**Figure 5:**
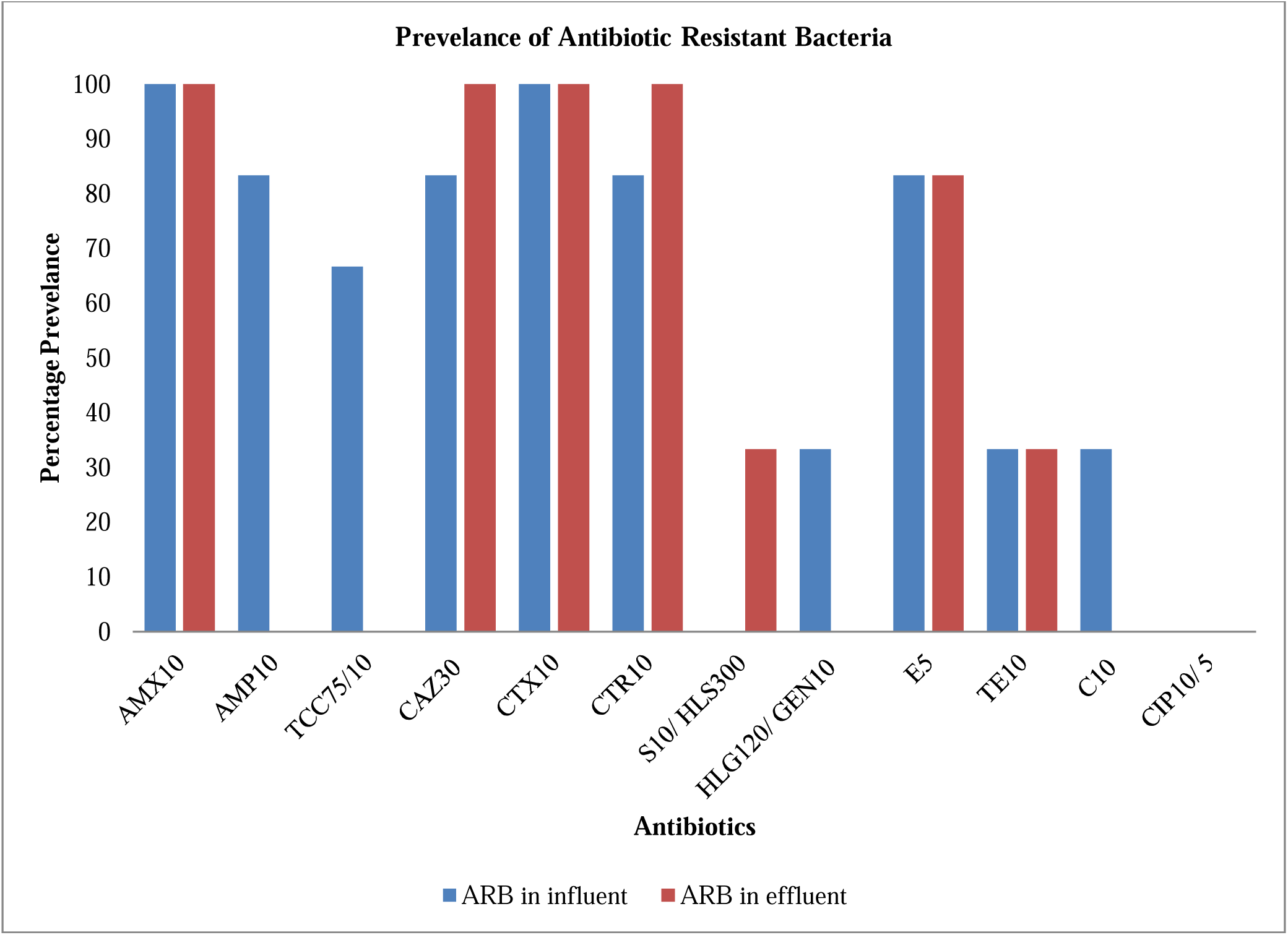
Prevalence of antibiotic resistant bacteria in influent and effluent

### 3.4 Molecular profiling of ARGs for ESBL, MRSA and Colistin resistant strains

Molecular characterization of different antimicrobial resistance bacteria was done by the presence or absence of various molecular markers like bla_SHV_, bla_TEM_ and bla_CTX-M_ genes for ESBL, *mecA* gene for MRSA and *mcr*-1 gene for Colistin in the effluent samples from the VF. The sample was screened for the presence of such marker genes by PCR using the primer against these genes markers. The results show that only bla_CTX-M_ gene with a band size of 544 bp was amplified indicating the presence of resistance against ESBL drug group Cephalosporin. However, all other resistant gene markers were not present in the effluent sample. MRSA isolates, having *mec*A gene are reported to be resistant to penicillin and oxytetracycline whereas no resistance was observed against ciprofloxacin, chloramphenicol and gentamicin [34]. This corroborated the antibiotic susceptibility test and even further validated the decrease in resistance in the non culturable species in the effluent samples as well. The production of ESBLs is one of the most important mechanisms of antimicrobial resistance. Increase in the number of ESBL attributes to the emergence of bla_CTX-M_ producing enterobacteriaceae. It is reported that *E. coli* is the predominant CTX-M group I ESBL producing enterobacteriaceae and resistance to third generation cephalosporin (3GC) antibiotics is the possible reason for emergence and spread of CTX-M ESBL producing organisms [35]. This result correlates with the antibiotic susceptibility test as high resistance was shown against 3GCs.

*S. epidermidis* was found in the effluent which is believed to acts as a reservoir for *mecA* gene and transfer it to *S. aureus* by horizontal gene transfer [36]. There is a possibility that this species present in the effluent was sensitive, as methicillin sensitive *S. epidermis* (MSSE) [37], since, no bands were observed for *mecA* gene. However, it might also be possible that earthworm-microorganism interactions play a certain role in shifting the resistivity pattern. The *mcr* genes are responsible for horizontal transfer of colistin resistance. This gene is generally present in *Proteus* strains like *P. rettgeri, P. vulgaris and P. mirabilis* [38]. However, since *P. inconstans* was isolated from the VF effluent, *mcr* gene was likely to be absent.

## 4. Mechanisms elucidating the role of earthworms-microorganism’s interactions in paradigm shift in antibiotic resistance patterns

Vermifiltration is proven to be a natural and sustainable technology for the treatment of wastewater with combined action of earthworms and microorganisms. Pathogens form an inescapable component of wastewater and being a potential hazard to the well-being of humans, need to be removed from it. This study focused on the performance efficacy of VF plant situated at Dr. B. Lal Institute of Biotechnology for clinical laboratory wastewater. In this novel study, antibiotic resistance activity of bacterial strains isolated from the VF was analyzed. The main objective was to investigate the effect of earthworms on the resistant microbial community found in the clinical wastewater before and after the treatment. The antimicrobial resistance activity of isolated microbes against 12 different antibiotics was investigated by Disk Diffusion Method. It was observed that there is a decrease in overall percentage of drug resistant strains towards the different drugs observed in the VF treated effluent as compared to the influent. This can be explained by the combined action of earthworms and indigenous microbes present in the active layer of the VF. The mechanism illustrating the role of earthworms and microorganisms is depicted in Figure 6. High removal percentages detected in the VF effluent is influenced by various microbial communities [43]. As a result, earthworms and indigenous microbes together form a biofilm and upon exposure to this biofilm, the ARB presents in the influent experiences a shift in the antibiotic resistivity pattern. It has been reported that earthworms could effectively decline tetracycline and fluoroquinolone resistance genes in activated sludge system as compared to the stabilization sludge system and without using earthworms [44]. The abundant bacteria in the earthworm gut make it a promising micro-zone for antibiotic degradation and degradation in soil is mainly carried out by *Pseudomonas, Staphylococcus*, and *Planctomycetes* degradation [45]. Therefore, the abundance of these bacteria strains was significantly increased in the worm gut or the adjacent soil when exposed to antibiotics and antibiotic resistant bacteria. Thus, this shift in the paradigm of wastewater might be due to long term exposure of ARB to the biofilm formed as a result of the interaction between earthworms and indigenous microbes. Yet more research is needed in the future to clarify the detailed information about the degradation mechanism of worm intestinal bacteria on ARB.

**Figure 6:**
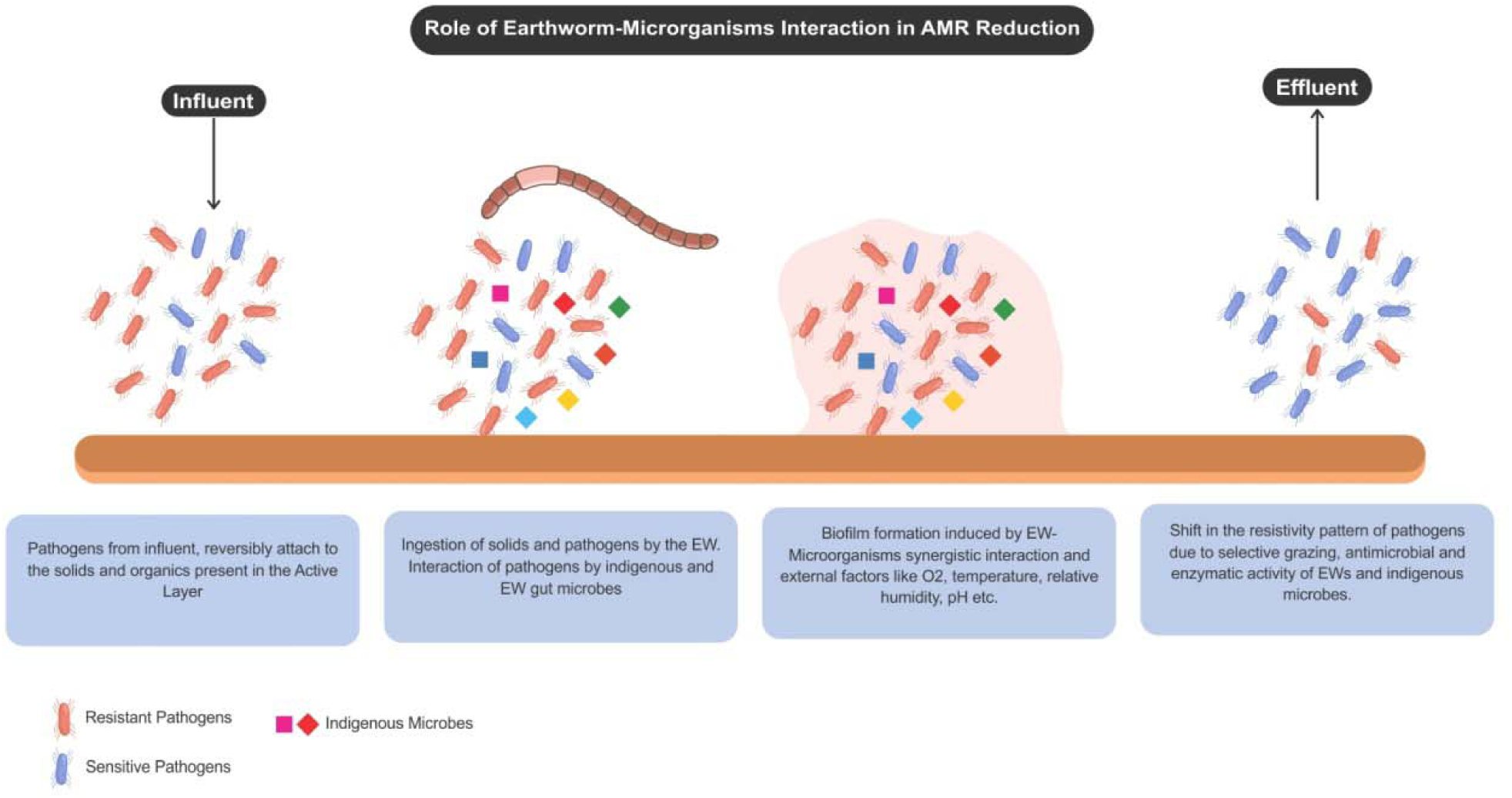
Role of earthworm-microorganisms interactions in biofilm formation and reduction of AMR from clinical laboratory wastewater

Earthworms represent the dominant invertebrate in soil and promote the decomposition of organic pollutants. They are one of the typical saprozoic organisms, living in the environment replete with microorganisms some of which may be a threat to their existence. It has been reported that to survive in such an environment, they have developed efficient immune-defense mechanisms against invading microorganisms [39]. Antibacterial peptides present in earthworms are listed as an important defense component against antimicrobial resistance. Several anti-infection components such as lysozymes and fetidins have been found in the earthworms[40].

The presence of earthworms can also stimulate microbial activities and population, which in turn accelerate the degradation of pollutants present in wastewater [41]. As described in Figure 7, VF proved to be an excellent artificial system through the inoculation of earthworms into a conventional biofilter which works on the principle of earthworm-microorganism symbiotic and synergistic interactions to treat clinical laboratory wastewater. Earthworm bioturbation impacts on the fate of contaminants by modifying their mobility and sorption to soil, leading to the change of the quality and dynamic of soil organic materials. They also provide and generate hot spots of microbial activity. Additionally, they perform effective degradation and transformation directly, through generating enzymes and, indirectly, owing to the microflora present in their intestine [42,43]. The fate of antibiotic resistance genes during six different full-scale municipal wastewater treatment processes and revealed that horizontal and vertical gene transfer mainly occurs in aerated tanks whereas anaerobic or anoxic tanks can reduce the occurrence of antibiotic resistance [46]. But to evaluate whether vertical and horizontal gene transfer is the major process for dissemination of antibiotic resistance within VF, a deeper analysis is required. An intriguing observation was the improvement in the overall activity of cell wall targeting and protein synthesis inhibiting drugs. Several studies have reported the presence of selective foraging in earthworms for solids, organic particles [47], microorganisms and other pathogens [21,48], however, further analysis might clearly reveal their specificity towards ARB.

**Figure 7:**
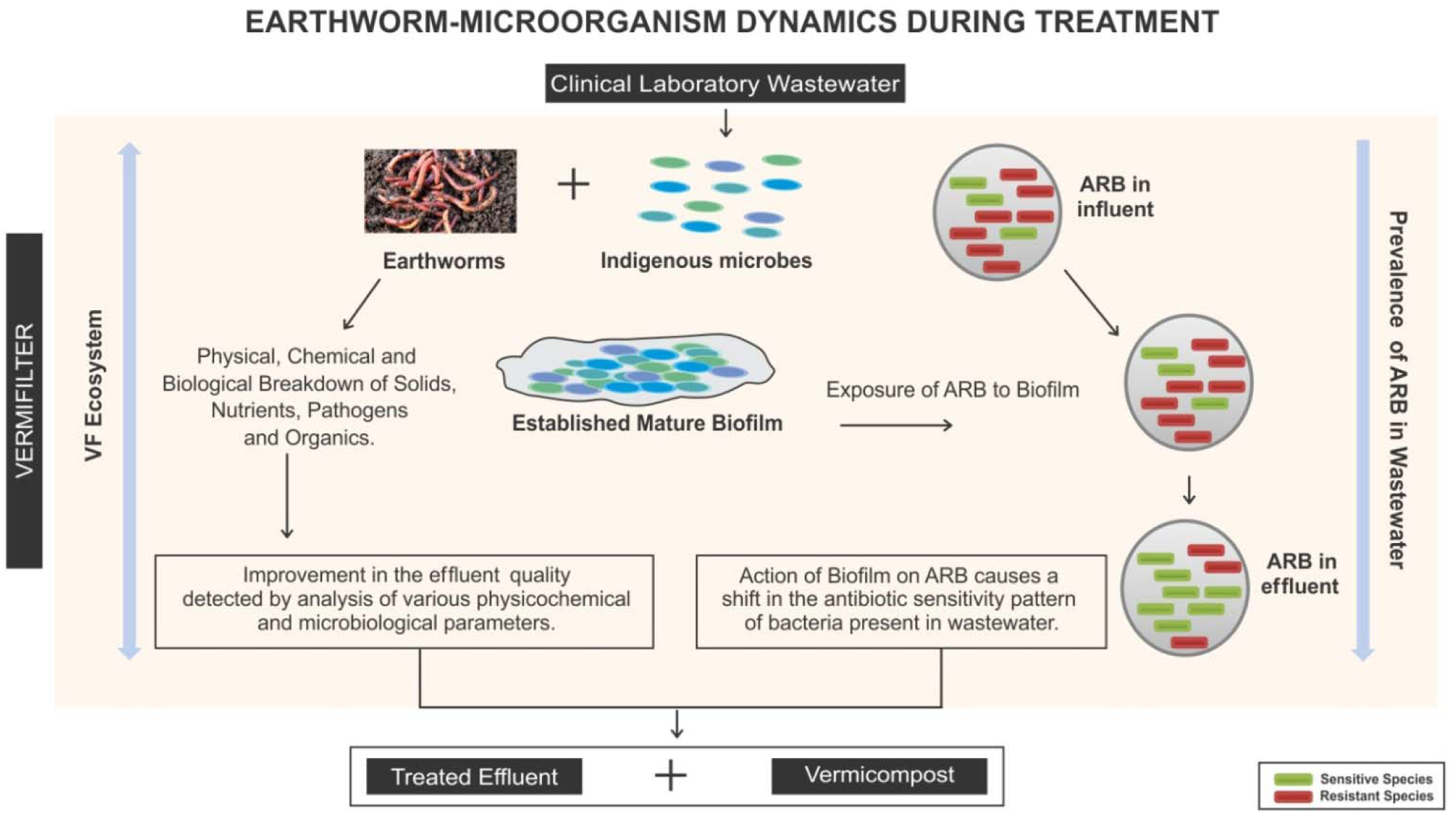
Dynamics involved in the treatment of clinical laboratory wastewater

It should also be noted that the presence of antibiotic compounds is not the only factor influencing the fate of antibiotic resistance. Therefore, other factors should also be taken into account such as the operational parameters of VF, the wastewater-associated bacterial communities and their interaction with other microbial communities, which may contribute to the spread of resistance genes associated with certain families of microorganisms. Conclusively, VF technology offers a sustainable solution for clinical wastewater with reliable performance in terms of reduction of physicochemical and microbiological parameters for clinical wastewater treatment. Also, since the effluents of hospitals contain high quantities of various ARBs, vermifiltration could be considered as a promising and economical technology for tertiary or refining treatment of these kinds of wastewater[49].

It should be recalled that this study focused on the ARB present in the wastewater fraction; however, further studies alluding to antibiotics and antibiotic resistance genes might provide a consolidated result helping in understanding the relationship between earthworm-microorganism activity and ARB. Despite this confinement, the combined culture-based and molecular approach used in this study provides a deeper understanding of the antimicrobial resistance community present in clinical laboratory wastewater.

## 5. Conclusions

On-site treatment of clinical laboratory wastewater is vital for the health and safety of surface waters used for recreation, drinking water, and animal habitat. The results confirmed the potential of vermifiltration technology for the treatment of clinical laboratory wastewater. Also, the study focused on selecting the potential bacterial strains from influent and effluent that possess antimicrobial resistance. It was found that earthworm-microorganism interactions play a crucial role in the regulation of physicochemical and microbiological parameters and additionally was responsible for causing a shift in the antimicrobial resistance pattern by the help of biofilm formation. Most efficient activity of antimicrobial resistance was found against drugs targeting cell wall and inhibiting protein synthesis. Together, these results suggest that vermifilter prove to be natural and sustainable on-site wastewater treatment technology for clinical laboratory wastewater thus, reducing the dissemination and proliferation of pathogens and ARB entering into the water streams.

## Credit authorship contribution statement

**Sudipti Arora (Corresponding author)**: Conceptualization, Investigation, Resources, Manuscript Writing and editing, corresponding author

**Sakshi Saraswat:** Experimental design, protocol standardization, Experimental conduction, Data analysis, Writing -original draft of manuscript.

**Ankur Rajpal**: Manuscript review

**Harshita Shringi:** Sampling, Experimentation, Data analysis

**Rinki Mishra:** Experimental design, protocol standardization, Experimental conduction, Data analysis

**Jasmine Sethi:** Sampling, Experimentation, Data analysis

**Jayana Rajvanshi**: Sampling, Experimentation, Data analysis

**Aditi Nag**: Experimental design on molecular work, protocol standardization, Experimental conduction

**Sonika Saxena:** Resources providing, Sample collection protocol standardization and supervision

**A. A. Kazmi:** Conceptualization and manuscript review

## Acknowledgements

The study group would like to acknowledge the constant support received from Dr. B. Lal Gupta (Director, Dr. B. Lal Institute of Biotechnology, Jaipur) and Dr. Aparna Datta for inspiring this research and providing daily motivations to work faster. The team further acknowledges the efforts made by research students Ms. Vikky Sinha, Ms. Vaishali Saboo and Mr. Harshit Chhabra for conducting experiments.

## References

[1] Kumar M, Jaiswal S, Sodhi KK, Shree P, Singh DK, Agrawal PK, et al. Antibiotics bioremediation: Perspectives on its ecotoxicity and resistance. Environ Int 2019. https://doi.org/10.1016/j.envint.2018.12.065.

[2] García J, García-Galán MJ, Day JW, Boopathy R, White JR, Wallace S, et al. A review of emerging organic contaminants (EOCs), antibiotic resistant bacteria (ARB), and antibiotic resistance genes (ARGs) in the environment: Increasing removal with wetlands and reducing environmental impacts. Bioresour Technol 2020. https://doi.org/10.1016/j.biortech.2020.123228.

[3] Momeni M. Hospital wastewater treatment scenario around the globe 2020;21:1–9.

[4] Beattie RE, Skwor T, Hristova KR. Survivor microbial populations in post-chlorinated wastewater are strongly associated with untreated hospital sewage and include ceftazidime and meropenem resistant populations. Sci Total Environ 2020. https://doi.org/10.1016/j.scitotenv.2020.140186.

[5] Krarup J, Jakob S, Nielsen U, Stefan Z. New standard for hospital wastewater treatment. Filtr + Sep 2015. https://doi.org/10.1016/s0015-1882(15)30141-5.

[6] Al-Enazi MS. Evaluation of Wastewater Discharge from Al-Sadr Teaching c Al-Khorah channel and Shatt Al-Arab River in Basra City-Iraq. J Environ Earth Sci 2016.

[7] Safe management of wastes from health care activities. Bull World Health Organ 2001. https://doi.org/10.1590/S0042-96862001000200013.

[8] CPCB. Environmental Standards for Ambient Air, Automobiles, Fuels, Industries and Noise 2001.

[9] Fouz N, Pangesti KNA, Yasir M, Al□Malki AL, Azhar EI, Hill□Cawthorne GA, et al. The contribution of wastewater to the transmission of antimicrobial resistance in the environment: Implications of mass gathering settings. Trop Med Infect Dis 2020;5. https://doi.org/10.3390/tropicalmed5010033.

[10] Salcedo DE, Lee JH, Ha UH, Kim SP. The effects of antibiotics on the biofilm formation and antibiotic resistance gene transfer. Desalin Water Treat 2015;54:3582–8. https://doi.org/10.1080/19443994.2014.923206.

[11] Gullberg E, Cao S, Berg OG, Ilbäck C, Sandegren L, Hughes D, et al. Selection of resistant bacteria at very low antibiotic concentrations. PLoS Pathog 2011;7:1–9. https://doi.org/10.1371/journal.ppat.1002158.

[12] Ragan MA, Beiko RG. Lateral genetic transfer: Open issues. Philos Trans R Soc B Biol Sci 2009;364:2241–51. https://doi.org/10.1098/rstb.2009.0031.

[13] Wang J, Chu L, Wojnárovits L, Takács E. Occurrence and fate of antibiotics, antibiotic resistant genes (ARGs) and antibiotic resistant bacteria (ARB) in municipal wastewater treatment plant: An overview. Sci Total Environ 2020;744:140997. https://doi.org/10.1016/j.scitotenv.2020.140997.

[14] Laht M, Karkman A, Voolaid V, Ritz C, Tenson T, Virta M, et al. Abundances of tetracycline, sulphonamide and beta-lactam antibiotic resistance genes in conventional wastewater treatment plants (WWTPs) with different waste load. PLoS One 2014;9:1–8. https://doi.org/10.1371/journal.pone.0103705.

[15] Bueno I, Williams-Nguyen J, Hwang H, Sargeant JM, Nault AJ, Singer RS. Systematic Review: Impact of point sources on antibiotic-resistant bacteria in the natural environment. Zoonoses Public Health 2018. https://doi.org/10.1111/zph.12426.

[16] Lin H, Zhang J, Chen H, Wang J, Sun W, Zhang X, et al. Effect of temperature on sulfonamide antibiotics degradation, and on antibiotic resistance determinants and hosts in animal manures. Sci Total Environ 2017. https://doi.org/10.1016/j.scitotenv.2017.07.057.

[17] Arora S, Rajpal A, Bhargava R, Pruthi V, Bhatia A, Kazmi AA. Antibacterial and enzymatic activity of microbial community during wastewater treatment by pilot scale vermifiltration system. Bioresour Technol 2014. https://doi.org/10.1016/j.biortech.2014.05.041.

[18] Li X, Yang J, Zhao L, Xing M, Deng D, Yi D. Profiling of Microbial Community Diversity in Vermifiltration of Different Earthworm abundances by PCR-DGGE 2009:25–7.

[19] Dhadse S, Satyanarayan S, Chaudhari PR, Wate SR. Vermifilters: A tool for aerobic biological treatment of herbal pharmaceutical wastewater. Water Sci Technol 2010. https://doi.org/10.2166/wst.2010.523.

[20] Ghatnekar et al. 2010. Application of Vermi-filter-based Effluent Treatment Plant (Pilot scale) for Biomanagement of Liquid Effluents from the Gelatine Industry. Dyn Soil, Dyn Plant 2010.

[21] Sinha RK, Bharambe G, Chaudhari U. Sewage treatment by vermifiltration with synchronous treatment of sludge by earthworms: A low-cost sustainable technology over conventional systems with potential for decentralization. Environmentalist 2008. https://doi.org/10.1007/s10669-008-9162-8.

[22] Arora S, Saraswat S, Mishra R, Rajvanshi J, Sethi J, Nag A, et al. Design, Performance Evaluation and Investigation of the Dynamic Mechanisms of Earthworm-Microorganisms interactions for wastewater treatment through Vermifiltration technology 2020;21:1–9. https://doi.org/10.1101/2020.08.14.252072.

[23] Bergey’s Manual® of Systematic Bacteriology. 2010. https://doi.org/10.1007/978-0-387-68572-4.

[24] Hudzicki J. Kirby-Bauer Disk Diffusion Susceptibility Test Protocol Author Information. Am Soc Microbiol 2012:1–13.

[25] Clinical and Laboratory Standards Institute(CLSI). Performance Standards for Antimicrobial Susceptibility Testing; 27th edition. CLSI supplement M100. 2017.

[26] Drieux L, Brossier F, Sougakoff W, Jarlier V. Phenotypic detection of extended-spectrum β-lactamase production in Enterobacteriaceae: Review and bench guide. Clin Microbiol Infect 2008. https://doi.org/10.1111/j.1469-0691.2007.01846.x.

[27] Arora S, Rajpal A, Kazmi AA. Antimicrobial Activity of Bacterial Community for Removal of Pathogens during Vermifiltration. J Environ Eng (United States) 2016. https://doi.org/10.1061/(ASCE)EE.1943-7870.0001080.

[28] Wang L, Zheng Z, Luo X, Zhang J. Performance and mechanisms of a microbial-earthworm ecofilter for removing organic matter and nitrogen from synthetic domestic wastewater. J Hazard Mater 2011. https://doi.org/10.1016/j.jhazmat.2011.08.035.

[29] Singh R, Samal K, Dash RR, Bhunia P. Vermifiltration as a sustainable natural treatment technology for the treatment and reuse of wastewater: A review. J Environ Manage 2019;247:140–51. https://doi.org/10.1016/j.jenvman.2019.06.075.

[30] Liu J, Lu Z, Yang J, Xing M, Yu F, Guo M. Effect of earthworms on the performance and microbial communities of excess sludge treatment process in vermifilter. Bioresour Technol 2012. https://doi.org/10.1016/j.biortech.2012.04.096.

[31] Liu G, Wang J. Long-term low DO enriches and shifts nitrifier community in activated sludge. Environ Sci Technol 2013. https://doi.org/10.1021/es304647y.

[32] Lederman ER, Crum NF. Pyogenic liver abscess with a focus on Klebsiella pneumoniae as a primary pathogen: An emerging disease with unique clinical characteristics. Am J Gastroenterol 2005. https://doi.org/10.1111/j.1572-0241.2005.40310.x.

[33] B Khyade V. Bacterial diversity in the alimentary canal of earthworms. J Bacteriol Mycol Open Access 2018. https://doi.org/10.15406/jbmoa.2018.06.00200.

[34] Kumar A, Kaushik P Anjay, Kumar P, Kumar M. Prevalence of methicillin-resistant Staphylococcus aureus skin and nasal carriage isolates from bovines and its antibiogram. Vet World 2017. https://doi.org/10.14202/vetworld.2017.593-597.

[35] Priyadharsini RI, Kavitha A, Rajan R, Mathavi S, Rajesh KR. Prevalence of bla CTX M Extended Spectrum Beta Lactamase Gene in Enterobacteriaceae from Critical Care Patients. J Lab Physicians 2011;3:080–3. https://doi.org/10.4103/0974-2727.86838.

[36] Najar-Peerayeh S, Moghadas AJ, Behmanesh M. Antibiotic susceptibility and mecA frequency in staphylococcus epidermidis, isolated from intensive care unit patients. Jundishapur J Microbiol 2014. https://doi.org/10.5812/jjm.11188.

[37] Bhimraj A. Postoperative Neurosurgical Infections. Elsevier Inc.; 2018. https://doi.org/10.1016/B978-0-323-32106-8.00050-9.

[38] Aghapour Z, Gholizadeh P, Ganbarov K, Bialvaei AZ, Mahmood SS, Tanomand A, et al. Molecular mechanisms related to colistin resistance in enterobacteriaceae. Infect Drug Resist 2019;12:965–75. https://doi.org/10.2147/IDR.S199844.

[39] Riera Romo M, Pérez-Martínez D, Castillo Ferrer C. Innate immunity in vertebrates: An overview. Immunology 2016. https://doi.org/10.1111/imm.12597.

[40] Wang X, Wang X, Zhang Y, Qu X, Yang S. An antimicrobial peptide of the earthworm Pheretima tschiliensis: cDNA cloning, expression and immunolocalization. Biotechnol Lett 2003. https://doi.org/10.1023/A:1024999206117.

[41] Wang Y, Xing M, Yang J. Earthworm (Eisenia fetida) Eco-physiological Characteristics in Vermifiltration System for Wastewater Treatment Through Analyzing Differential Proteins. Water Air Soil Pollut 2017. https://doi.org/10.1007/s11270-016-3138-y.

[42] Mougin C, Cheviron N, Repincay C, Hedde M, Hernandez-Raquet G. Earthworms highly increase ciprofloxacin mineralization in soils. Environ Chem Lett 2013. https://doi.org/10.1007/s10311-012-0385-z.

[43] Ghobadi N, Shokoohi R, Rahmani AR, Samadi MT, Godini K, Samarghandi MR. Performance of A Pilot-Scale Vermifilter for the Treatment of A Real Hospital Wastewater. Avicenna J Environ Heal Eng 2016. https://doi.org/10.5812/ajehe-7585.

[44] Huang K, Xia H, Wu Y, Chen J, Cui G, Li F, et al. Effects of earthworms on the fate of tetracycline and fluoroquinolone resistance genes of sewage sludge during vermicomposting. Bioresour Technol 2018. https://doi.org/10.1016/j.biortech.2018.03.021.

[45] Sun M, Chao H, Zheng X, Deng S, Ye M, Hu F. Ecological role of earthworm intestinal bacteria in terrestrial environments: A review. Sci Total Environ 2020. https://doi.org/10.1016/j.scitotenv.2020.140008.

[46] Tong J, Tang A, Wang H, Liu X, Huang Z, Wang Z, et al. Microbial community evolution and fate of antibiotic resistance genes along six different full-scale municipal wastewater treatment processes. Bioresour Technol 2019. https://doi.org/10.1016/j.biortech.2018.10.079.

[47] Bhadauria T, Saxena KG. Role of Earthworms in Soil Fertility Maintenance through the Production of Biogenic Structures. Appl Environ Soil Sci 2010. https://doi.org/10.1155/2010/816073.

[48] Kale RD, Karmegam N. The Role of Earthworms in Tropics with Emphasis on Indian Ecosystems. Appl Environ Soil Sci 2010. https://doi.org/10.1155/2010/414356.

[49] Tehrani AH, Gilbride KA. A closer look at the antibiotic-resistant bacterial community found in urban wastewater treatment systems. Microbiologyopen 2018. https://doi.org/10.1002/mbo3.589.

